# Structural basis for substrate recognition, ligation and activation by a hyperactive Asn peptide ligase from *Viola yedoensis*

**DOI:** 10.1101/2021.12.09.471967

**Authors:** Side Hu, Abbas El Sahili, Srujana Kishore, Yee Hwa Wong, Xinya Hemu, Boon Chong Goh, Zhen Wang, James P. Tam, Chuan-Fa Liu, Julien Lescar

## Abstract

Peptide asparaginyl ligases (PALs) belong to a limited class of enzymes from cyclotide-producing plants, that perform site-specific ligation reactions after a target peptide Asx (Asn/Asp) binds to the ligase active site. How PALs specifically recognize their polypeptide substrates has remained elusive especially at the prime binding side of the enzyme. Here we captured VyPAL2, a catalytically efficient PAL from *Viola yedoensis*, in an activated state, with and without a bound substrate. The bound structure shows one ligase with the N-terminal polypeptide tail from another ligase molecule trapped at its active site, revealing how Asx inserts in the enzyme’s S1 pocket and why a hydrophobic residue is required at the substrate P2’ position. Beside illustrating the role played by P1 and P2’ residues as primary anchors for the enzyme reaction, these results provide a mechanistic explanation for the role of the “Gatekeeper” residue at the surface of the S2 pocket, in shifting the non-prime portion of the substrate and, as a result, the activity towards either ligation or hydrolysis. These results detail the molecular events that occur during proenzyme maturation in the plant vacuolar compartment, suggest a mechanism for ligation, and will inform the design of peptide ligases with tailored specificities.

**One sentence summary:** We captured VyPAL2, a catalytically efficient plant peptide ligase with a bound substrate, providing the molecular basis for substrate recognition and ligation.

## INTRODUCTION

Peptide ligases, such as the Sortase A (Schneewind et al., 1992), constitute attractive tools for protein conjugation (Mao et al., 2004), protein semi-synthesis (Noike et al., 2015; Cao et al., 2016), peptide cyclization (Antos et al., 2009) or live cell labelling (Bi et al., 2017). Ligases extracted from plants could play a crucial role in the active field of “molecular epigenetics” where precision tools are needed for instance to produce exquisitely modified histone proteins (Bagert and Muir, 2021). Following the discovery of cyclotide-like protein sequences in graminaceous crop plants (Mulvenna et al., 2006) and the realization that these cyclic polypeptides depended on plant asparaginyl endopeptidases for maturation (Mylne et al., 2012), a quest to clone these enzymes was started. In 2014, a peptide ligase, specific for Asn, was successfully isolated from the cyclotide-producing plant Clitoria ternatea and named butelase 1 (Nguyen et al., 2014). Compared to the bacterial Sortase A, butelase 1 is a hyperactive peptide ligase (PAL) with a reported catalytic efficiency of 1,314,000 M^-1^s^-1^ that has proven useful for many applications (Cao et al., 2016; Bi et al., 2017; Nguyen et al., 2015a, 2015b; Cao et al., 2015; Nguyen et al., 2016a, 2016b). Subsequently, several PALs including OaAEP1b (Harris et al., 2015), HeAEP3 (Jackson et al., 2018) and VyPAL2 (Hemu et al., 2019), were identified from other cyclotide-producing plants: Oldenlandia affinis, Hybanthus enneaspermus, and Viola yedoensis respectively.

Asparaginyl endopeptidases (AEPs or legumains, EC 3.4.22.34 (Kembhavi et al., 1993)) belong to the subfamily C13 of cysteine protease. Legumains such as butelase 2 (Serra et al., 2016), OaAEP2 (Serra et al., 2016) and HaAEP1 from the sunflower Helianthus annuus (Haywood et al., 2018) are predominantly endowed with protease activity at neutral pH, with a low level of peptide ligase activity (Haywood et al., 2018; Dall and Brandstetter, 2013; Zhao et al., 2014; Zauner et al., 2018a; Dall et al., 2020, 2021). A third group of enzymes, including CeAEP from jack bean (Bernath-Levin et al., 2015), PxAEP3 from Petunia x hybrida (Jackson et al., 2018), AtLEGγ from Arabidopsis thaliana (Zauner et al., 2018a) or VyAEP1 from Viola yedoensis (Hemu et al., 2019), catalyse, at near neutral pH, the formation of both ligation and hydrolytic products from peptide substrates carrying proper amino acid recognition sequences. In contrast to these two groups of “bi-functional” or “predominant” AEPs, butelase 1 (Nguyen et al., 2014), OaAEP1b (Harris et al., 2015), OaAEP3-5 (Harris et al., 2019) and VyPAL2 (Hemu et al., 2019) mostly catalyze the formation of ligation products, whilst little to no hydrolytic product is observed at either neutral or acidic pH.

In plants, all these enzymes, (having either predominantly Asx peptide ligase, endoprotease or a hybrid protease/ligase activity), are expressed as inactive zymogens. The expressed protein contains a vacuolar addressing signal peptide, an N-terminal pro-domain, a catalytic core domain and a C-terminal cap domain that covers the active site of the folded pro-enzyme (Kuroyanagi et al., 2002). Auto-activation is performed in vivo in the acidic vacuolar compartment of the plant, leading to the conversion of the pro-enzyme to its mature active form, via proteolysis of the immature polypeptide chain presumably in trans (Nguyen et al., 2014). Similarly, in vitro activation of the recombinant proenzyme is usually performed in the laboratory at pH values ranging from 4 to 5: under these acidic conditions, the zymogen undergoes autolytic activation leading to the removal of both the pro- and cap domains at the N- and C-termini of the catalytic core (Harris et al., 2015; Jackson et al., 2018; Hemu et al., 2019; Yang et al., 2017) and to the release of mature active enzymes with an active site fully exposed to the solvent. Auto-activation is thus the one-off capacity of these enzymes to catalyse a hydrolysis reaction, regardless of the dominant activity of the activated mature form (protease or ligase) raising the question of the precise catalytic mechanism underlying enzyme auto-activation.

Both AEPs and PALs share a high level of structural similarity, suggesting that their enzymatic activity are controlled by subtle structural differences at key positions near the catalytic center (Hemu et al., 2019). Recent structure-function studies identified important determinants for peptide formation, which were named “marker of ligase activity” (Jackson et al., 2018) and “Ligase activity determinants” (Hemu et al., 2019). Local structural features, immediately surrounding the S1 Asx binding site, were proposed to control (i) the conformation of the bound peptide substrates (ii) the kinetics of peptide departure at the enzyme non-prime site (iii) the access to the S-acyl enzyme intermediate of either water molecules, leading to hydrolysis of the peptide substrate, or alternatively of an incoming nucleophile - leading to ligation- at the prime side. Some empirical rules were proposed (Hemu et al., 2019), that were largely validated via the successful conversion of butelase 2, a pure Asx protease, into a “butelase 1-like” Asx peptide ligase by solely mutating residues of its S2 (Ligase Activity determinant 1 or LAD1) and S1’ (LAD2) substrate-binding pockets, while mutations at other non-LAD positions had only a limited impact on activity (Hemu et al., 2020).

Of specific functional importance is the “Gatekeeper” residue located at the surface of the S2 (LAD1) pocket. Initially discovered via a systematic site-directed mutagenesis study of OaAEP1b, a weakly-active PAL isolated from Oldenlandia affinis: the Gatekeeper which is Cys247 in the wild-type OaAEP1b enzyme, plays a key role in controlling both the directionality and the kinetics of the ligase vs protease reaction (Yang et al., 2017). A single mutation of Cys247 into Ala was sufficient to increase the value of kcat/Km for ligation by over 150-fold compared to the WT OaAEP1b enzyme. Likewise, the cognate substitution of Gatekeeper residue Gly252 of butelase 2 into residues such as Val or Ile, yielded butelase 2 mutants with a markedly increased ligase activity (Hemu et al., 2020). Thus, the presence of a Gly residue at the Gatekeeper position in the enzyme amino acid sequence is indicative of a predominantly protease activity for the mature enzyme, whilst residues with aliphatic side chains at this position favour ligase activity (Hemu et al., 2020, 2019).

Despite the publication of several plant AEP and PALs crystal structures, either in their pro-enzyme or active forms devoid of the cap domain (Hemu et al., 2019; Haywood et al., 2018; Dall and Brandstetter, 2013; Zhao et al., 2014; Zauner et al., 2018a; Yang et al., 2017; James et al., 2019), our understanding of the mechanistic and structural importance of LAD1 and LAD2 residues and of the Gatekeeper remains incomplete. Here we sought structural determinants accounting for substrate recognition and ligase activity of VyPAL2, a very efficient PAL discovered via mining the transcriptome of Viola yedoensis that can be conveniently expressed in a recombinant form (Hemu et al., 2019). We obtained crystal structures following acid-induced auto-activation, including one structure where residues from the N-terminal propeptide of one ligase molecule are bound to the active site of the neighbouring molecule in the crystal lattice. This structure gives for the time, to our knowledge, a complete atomic view of substrate binding including at the prime side of the enzyme. Together with supporting biochemical and enzymatic data, key structural features associated with specific substrate recognition are disclosed and a testable mechanism for peptide ligase activity is proposed in which the Gatekeeper plays a crucial role, which agrees with kinetics observations. Moreover, the study of auto-activation of VyPAL2 proenzymes bearing mutations at their His and Cys catalytic residues points to alternative routes used by these enzymes for their maturation process and for performing their catalytic activity.

## RESULTS AND DISCUSSION

### Ligase activity and crystal structure of the activated enzyme

VyPAL2 was expressed, purified, and activated using the reported procedure (Hemu et al., 2019). To conform to the LAD1 of butelase 1 which has the highest ligase activity documented so far (Nguyen et al., 2014; Hemu et al., 2019, 2020), a single mutant of the VyPAL2 proenzyme with the mutation I244V at the gatekeeper position (VyPAL2-I244V) was also expressed, purified, and similarly activated. Briefly, activation was carried out by proenzyme incubation at pH 4.4 for 2 hours at 37°C followed by size-exclusive chromatography to purify the core catalytic domain (Figure 1A; Supplemental Figure 1). Upon activation, for both VyPAL2 and VyPAL2-I244V, a time-dependent decrease in the amount of the 50 kDa pro-enzyme is observed on SDS-PAGE. This is accompanied by the apparition of a ∼37 kDa band corresponding to the active form and activation is completed after 2 hours incubation (Figures 1A and 1B).

**Figure 1.**
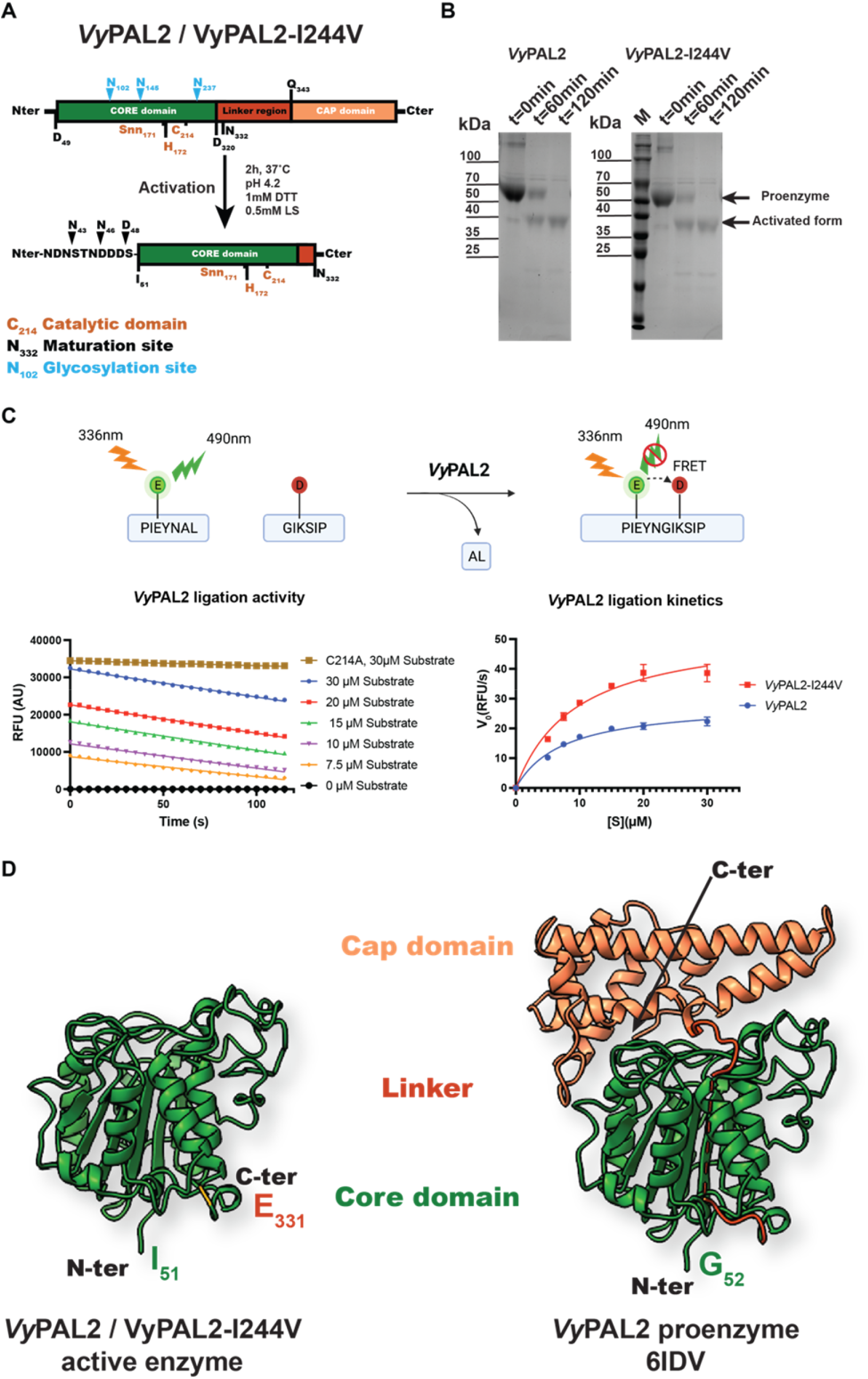
Expression, activation, ligase activity and crystal structure of VyPAL2-I244V. (A) Schematic sequence representation of VyPAL2-I244V. The respective VyPAL2 genes encoding complete amino acid sequences were cloned into the expression vector, with the signal peptide substituted by a hexa-His-tag for affinity purification (see methods). Catalytic residues and the aspartimide moiety (Snn171) next to the catalytic His172 are indicated in brown, N-linked glycosylation sites are indicated in cyan and domain boundaries as deduced from LC-MS/MS are indicated in black (please see also SI of ref 18). Proteolytic cleavage sites are indicated with arrows above the N- and C-terminal sequences. The products from low pH activation of the proenzyme were analysed using (B) SDS-PAGE with Coomassie blue staining. Incubation over two hours of VyPAL2 leads to a fully matured enzyme. (C) ligase activity using FRET assay. (D) Comparison between crystal structures of the VyPAL2 pro-enzyme monomer (right panel, PDB access code: 6IDV ref: 18) and the activated VyPAL2 protein (left panel, this work). The proteins are displayed with α-helices as ribbons and β-strands as arrows. The colour-code used for each of the three domains of the proenzymes is used throughout the manuscript (core domain: green, cap domain: wheat, linker region connecting the catalytic core and cap domain in orange).

Next, we determined the peptide ligase activity of the VyPAL2 WT enzyme and VyPAL2-I244V using a fluorometric assay allowing real-time monitoring of the formation of a ligated peptide product (Figure 1C). Two model peptide substrates with amino-acid sequences PIE(EDANS)YNAL and GIK(DABSYL)SIP, containing the “NAL” tripeptide and ‘GI” dipeptide ligase recognition motifs at their C- and N-termini respectively, were fluorescently labelled. Upon ligation, the fluorescence signal of the EDANS moiety of the first peptide becomes quenched by the DABSYL moiety of the second peptide, whilst the AL dipeptide is released (Figure 1C). Ligation reactions of VyPAL2 and VyPAL2-I244V were performed at 37 °C for 2 min to determine the initial velocity, at a pH of 6.5 (Figure 1C). The value of Vmax of VyPAL2-I244V (53.76 RFU/s) is increased by approximately 2-fold compared to the WT VyPAL2 enzyme (28.68 RFU/s), while the Km values were comparable with Km= 9.351 μM for the WT enzyme and 7.455 μM for VyPAL2-I244V (Figure 1C).

The purified auto-activated VyPAL2 and VyPAl2-I244V proteins were then concentrated to 6.7 mg/ml and 4.5 mg/ml respectively prior to crystallization. Well-diffracting crystals appearing after 3-5 days were mounted on a cryo-loop and flash frozen in liquid nitrogen. X-ray diffraction intensities extending to 2.3 and 1.59 Å resolution for VyPAL2 and VyPAl2-I244V respectively were collected and the structures of their active forms were determined by molecular replacement using the core domain of the VyPAL2 proenzyme structure as a search probe(Hemu et al., 2019) (Supplemental Table 1). Two essentially identical molecules are present in the asymmetric unit. The small interface between these two molecules rules out the formation of a functional dimer in solution coherent with the observed elution volume in size exclusion chromatography corresponding to a monomer with a ∼40 kDa apparent molecular weight. The structure of the active enzyme comprising residues Ile51 to Glu331 adopts the conserved core domain architecture also seen in legumains: a central β sheet containing six β strands surrounded by five large and four smaller α-helices (Figure 1D). Compared to the equivalent core domain in the context of the immature proenzyme VyPAL2 (Hemu et al., 2019), no major structural change occurs upon maturation, as indicated by a root mean square deviation (r.m.s.d) of 0.3 Å for 276 superimposed α-carbon (Figure 1D). Hence, activation appears to have little impact on the structure of the core domain, besides cap removal, leading to a complete exposure of the catalytic site to the solvent and to diffusing protein substrates (Figure 1D). The structures of VyPAL2 and VyPAL2-I224V are highly resembling with a r.m.s.d. of 0.171 Å (Supplemental Figure 4). Further comparisons with other active PALs such as AtLEGγ (Zauner et al., 2018b) (PDB access code: 5OBT), HaAEP1 (Haywood et al., 2018) (PDB code: 6AZT), or butelase 1 (James et al., 2019) (PDB code: 6DHI), return an average r.m.s.d. value of 0.5 Å (Supplemental Table 2) confirming that no major change of conformation occurs when enzymes undergo maturation.

To obtain further insights into the mechanism of auto-activation, we calculated the distribution of electrostatic charges at the interface of the VyPAL2 core with the cap domain (Supplemental Figure 2). Calculations were performed at a pH value of 6.5, where the proenzyme interface is stable, and at pH 4.4 near the pH found in the vacuole compartment and also used for in vitro activation of the proenzyme. At near neutral pH, residues exposed from the core domain are globally neutral or slightly negatively charged, while the contacting residues exposed at the surface of the cap domain bear a slightly positive charge (Supplemental Figure 2). Together with shape complementarity, these opposite electrostatic charges contribute to the formation of a stable interface between these two domains, explaining the large thermostability of the proenzyme. Upon acidification however, both the interface regions of the cap and core domains become strongly positively charged, due to the protonation of several His, Asp and Glu side chains exposed at the surface. This leads to a strong electrostatic repulsion between the core and cap domains, and interface destabilization is likely to facilitate access of the linker region to the enzyme active site favoring trans-activation and release of the mature core domain (Supplemental Movie 1).

**Figure 2.**
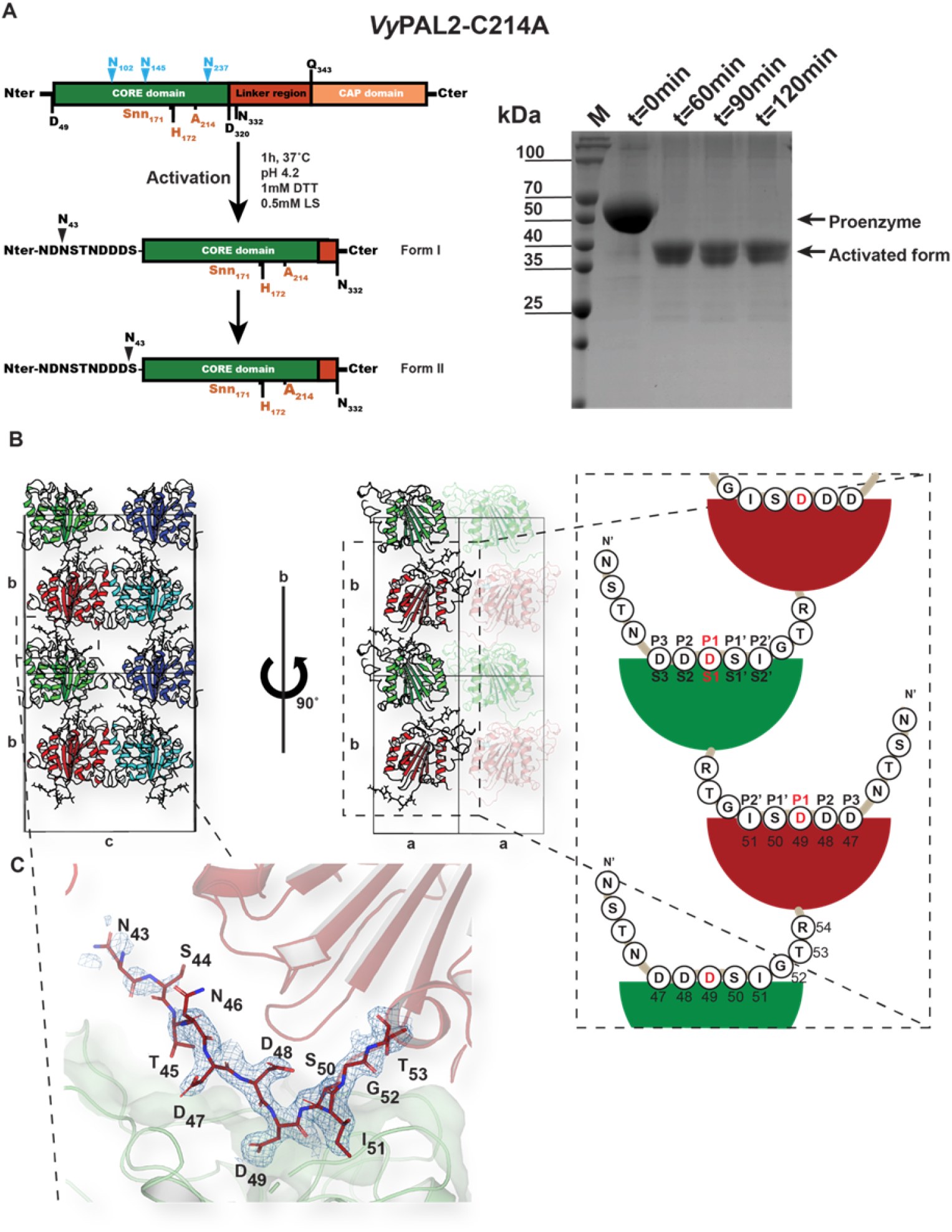
Expression, processing, and crystal structure of VyPAL2-C214A. (A) Schematic sequence representations of VyPAL2-C214A (left panel), and incubation over one hour of VyPAL2-C214A leads to a fully processed enzyme (“activated form”, right panel). (B) Schematic of the crystal packing showing how VyPAL2-C214A molecules are stacked in the monoclinic C2 unit cell. Molecules related by a unit-cell translation along b are displayed as ribbons with the same colour, whilst molecules related by the 2-fold axis along the same axis are displayed in green and red (middle panel). The N-terminal tail of each molecule inserts in the molecule stacked underneath. (C) The N-terminal peptide bound to the enzyme. The molecular surface of the peptide ligase is displayed in grey. Overlaid is a Fourier difference map displayed as a blue mesh with coefficients 2Fo-Fc with phases form the refined model, contoured at a level of 1σ over the mean.

### Crystallization of an enzyme-substrate complex

In order to trap a complex with a peptide substrate, a VyPAL2 single mutant was generated by mutating the catalytic Cys214 to alanine. Remarkably, this active site mutant undergoes rapid maturation at acidic pH and its core domain could be purified using a procedure similar to the wild-type enzyme (Figure 2A). Moreover, auto-activation of VyPAL2-C214A appears even faster than VyPAL2-I244V as full processing of the proenzyme is observed after only 60 minutes vs 2 hours for VyPAL2-I244V (Figures 1B and 2A; Supplemental Figure 3). Previous studies have critically examined the role of the oxyanion hole in the subtilisin protease (Bryan et al., 1986) and this will be discussed below in the light of the present data. As expected, VyPAL2-C214A showed a highly reduced activity in our peptide ligation assay, supporting a crucial role for Cys214 in the ligation and cyclization reactions but not for maturation.

**Figure 3.**
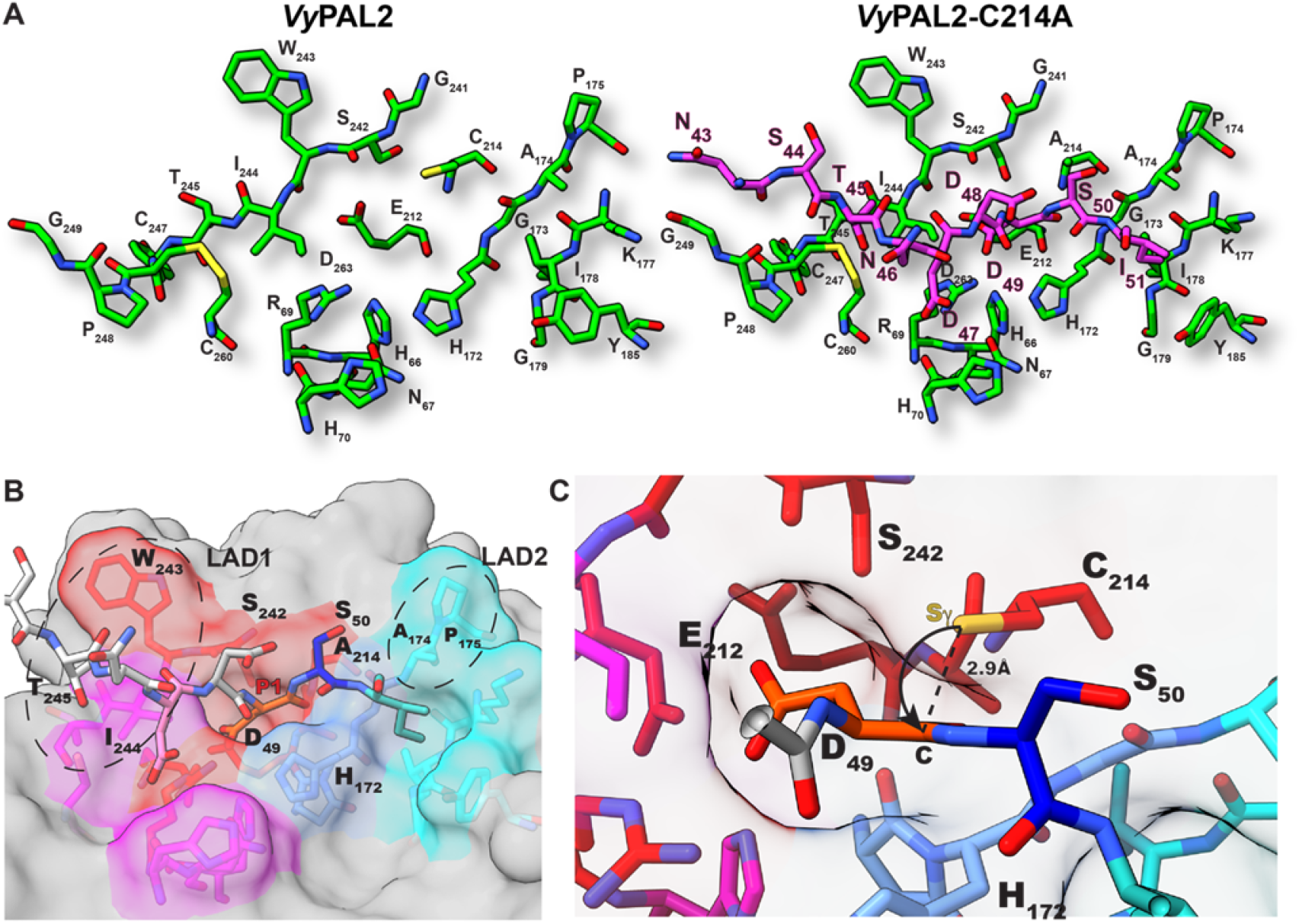
Peptide binding groove and substrate recognition by VyPAL2. (A) The left panel shows the peptide binding groove of the ligase as derived from the crystal structure of VyPAL2 (Supplemental Table 1, this work). The right panel is derived from the structure of VyPAL2-C214A (Supplemental Table 1, form I, this work) bound to the peptide displayed in magenta. (B) Residues lining the substrate binding pockets are depicted as sticks. VyPAL2 core domain is represented as a grey surface. The S1 pocket is red, the S3 pocket is magenta, the S1’ pocket is blue, S2’ pocket is cyan. (C) The active site residues His172 and C214 (mutated here in Ala) are on opposite sides of the peptide substrate and modelling back a sulfhydryl group at A214 shows that the distance with the scissile carbon is consistent with an inline attack of the carbon atom of the carbonyl group as displayed.

Following maturation at acidic pH and purification of the activated form, well-diffracting crystals of the core domain of VyPAL2-C214A could be obtained in two conditions (see methods): while form I crystals grow after 3-5 days, form II crystals only grow after 2 months incubation (Figure 2A). X-ray diffraction data at resolutions of 1.9 and 1.8 Å respectively could be collected for these two crystal forms (Supplemental Table 1). Form I crystals contain one monomer per asymmetric unit and molecules are packed in staggered rows along the unique b axis (Figure 2B). Residues Ser44 to Val329 of the protein, could be traced with confidence in the electron density map, including residues Asn43-Asp49 at the N-terminal cleavable proenzyme region which could be confidently built in well-defined electron density (Figure 2C). In particular, the scissile P1-P1’ (Schechter and Berger, 1967) bond Asp49-Ser50 is neither cleaved during the one-hour low pH activation step nor during the following purification and crystallization steps (over 3-5 days). As anticipated, this cleavable N-terminal tail appears flexible and projects away from the enzyme catalytic core. In the crystal lattice, this tail inserts into the active site of the neighbouring molecule related to the first by the crystallographic dyad along the b axis (Figures 2B and 2C). Thus, in the crystal lattice, the N-terminal region (Asn43-Asp49) of a molecule occupies the active sites of a molecule located immediately below (Figures 2B and 2C). Of note, Asp49 is deeply buried in the S1 pocket (Figures 3A and 3B) while residues Asp47, Asp48, Ser50 and Ile51 occupy the S3, S2, S1’ and S2’ pockets respectively (Figures 3 and 4). These observations agree with previously reported LC-MS/MS results demonstrating that the peptide bond C-terminal to Asp49 of VyPAL2 constitutes one possible N-terminal maturation site of the proenzyme18. Hence, the N-terminal peptide Asn43-Ile51 bound to the active site of the VyPAL2-C214A likely represents an enzyme-substrate complex trapped during N-terminal maturation. Beside the Asp49 cleavage site, several other possible proteolytic sites were uncovered using LC-MS/MS experiments at Asn43, Asn46 and Asp48 (see Figure S3 of Hemu et al., 2019).

**Figure 4.**
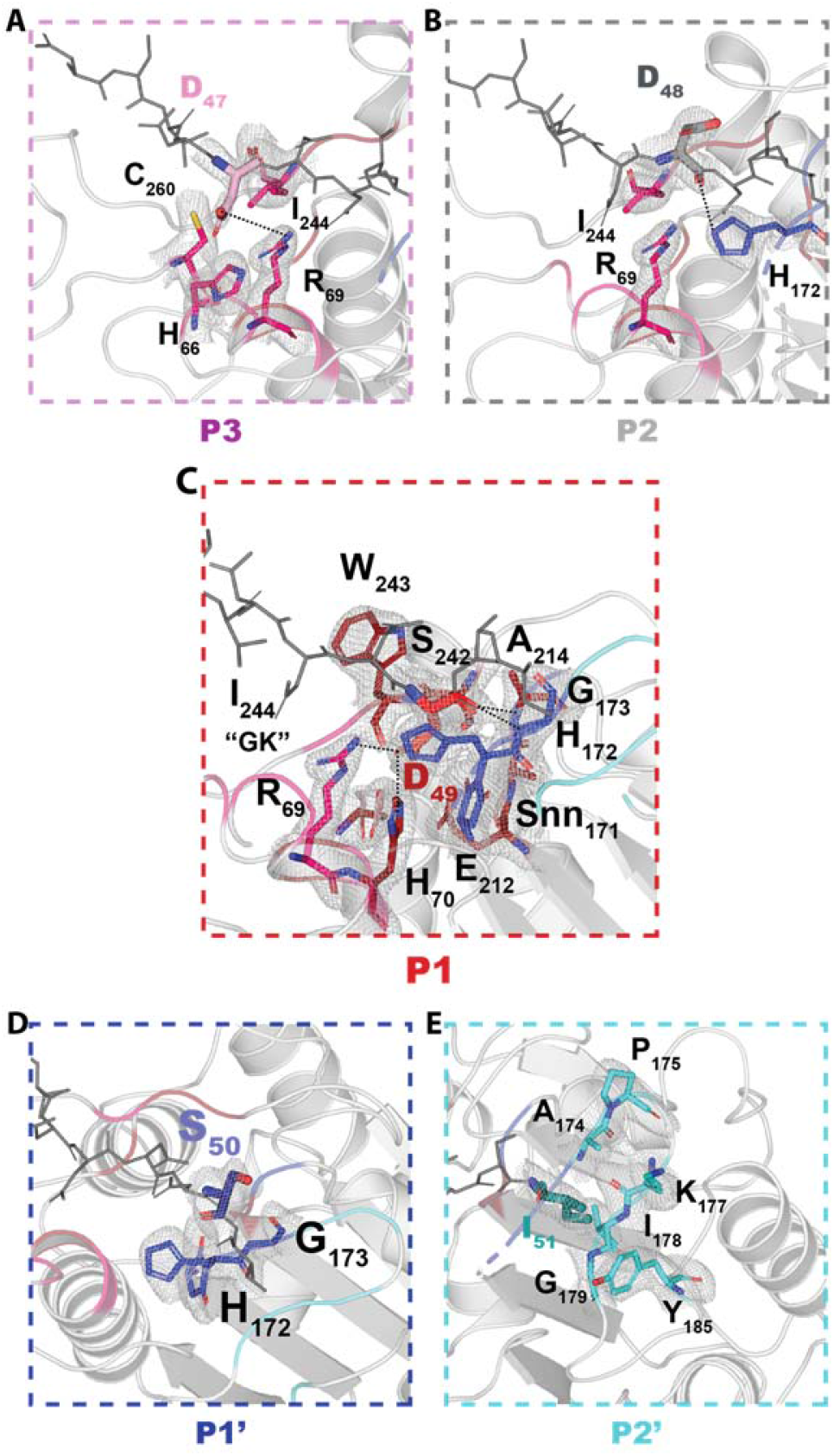
Detailed views of the ligase specificity pocket. Magnified views of the S3-S2’ specificity pockets of VyPAL2. For each pocket, the cognate peptide substrate (Sticks) is coloured in wheat (A, P3), grey (B, P2), red (C, P1) blue (D, P1’), and cyan (E, P2’) while the rest of the peptide is in grey. Residues from the specificity pockets are in coloured in magenta (A, S3), red (B, S2), red (C, S1) blue (D, S1’) and cyan (E, S2’) and shown as sticks and labelled. Polar interactions between the substrate residues and residues lining the specificity pockets (Hydrogen bonds or salt bridges) are depicted as dashed lines. Overlaid is a Fourier difference map displayed as a blue mesh with coefficients 2Fo-Fc with phases form the refined model, contoured at a level of 1σ over the mean.

Following low pH treatment, we obtained crystals of VyPAL2-C214A in another condition (form II; Supplemental Table 1). While form I crystals of VyPAL2-C214A with the bound N-terminal peptide described above (Supplemental Table 1) appeared within 5 days, form II crystals only grow after two-month incubation, an observation consistent with the much slower kinetics of auto-proteolysis of residues 43-49 by VyPAL2-C214A compared to the C-terminal cap region. In VyPAL2-C214A crystal form II, the N-terminal tail has been cleaved (Supplemental Figure 4) and the polypeptide chain now extends from Ser50 to Val329. Taken together, these results show that the VyPAL2-C214A complex structure (form I) represents an enzyme-substrate complex while form II captured the product of this slow N-terminal cleavage event.

### Overall architecture of the enzyme-substrate complex

Form I crystals offer a unique opportunity to analyse substrate binding for an activated PAL and to compare it to the unbound auto-activated structure. The bound N-terminal peptide sits in a shallow groove at the surface of the ligase and intermolecular interactions between the ligase and its substrate appears primarily mediated by P1 residue Asp49 and Ile51 (P2’), which act as two anchoring points (Figures 3A and 3B). The monomer can be superimposed with the activated VyPAL2-I244V structure with a r.m.s.d. 0.21 Å. Thus, binding of the Asn43-Ile51 N-terminal substrate is not accompanied with significant conformational changes (compare left and right panels in Figure 3A). Asp49 is bound in the S1 pocket as one would expect for the isosteric enzyme’s P1 substrate Asparagine. At acidic pH of 4.5 used to obtain form I crystals, both the carboxylic sidechain of Asp49 as well as of Glu212 and Asp263 present in the S1 pocket are likely to be protonated, favouring binding, while binding would be disfavoured at neutral pH (Zhang et al., 2021). Since the N-terminal Asn43-Ile51 region represents a catalytically functional substrate positioned in the active site, we used this experimental structure to simulate an active enzyme-substrate complex having a Cys back at position 214 bound to residues P3 to P2’. Such a complex allows the precise definition of the enzyme binding pockets S3-S2’ (Figures 3B and 4).

We then compared the mode of binding of this N-terminal peptide with the one observed when the cap binds to the same active site pocket. The proteolytically-sensitive segment from the cap plugged into the proenzyme active site, keeps it in an inactive state (Yang et al., 2017). Although the two polypeptide chains partially overlap especially the two amide groups from Gln323 and Asp49 in the S1 pocket, residues Asn43-Gly52 (this work) run in an opposite direction compared to residues Ile341-His344 of the cap domain (PDB access code: 6IDV; Supplemental Figure 5).

**Figure 5.**
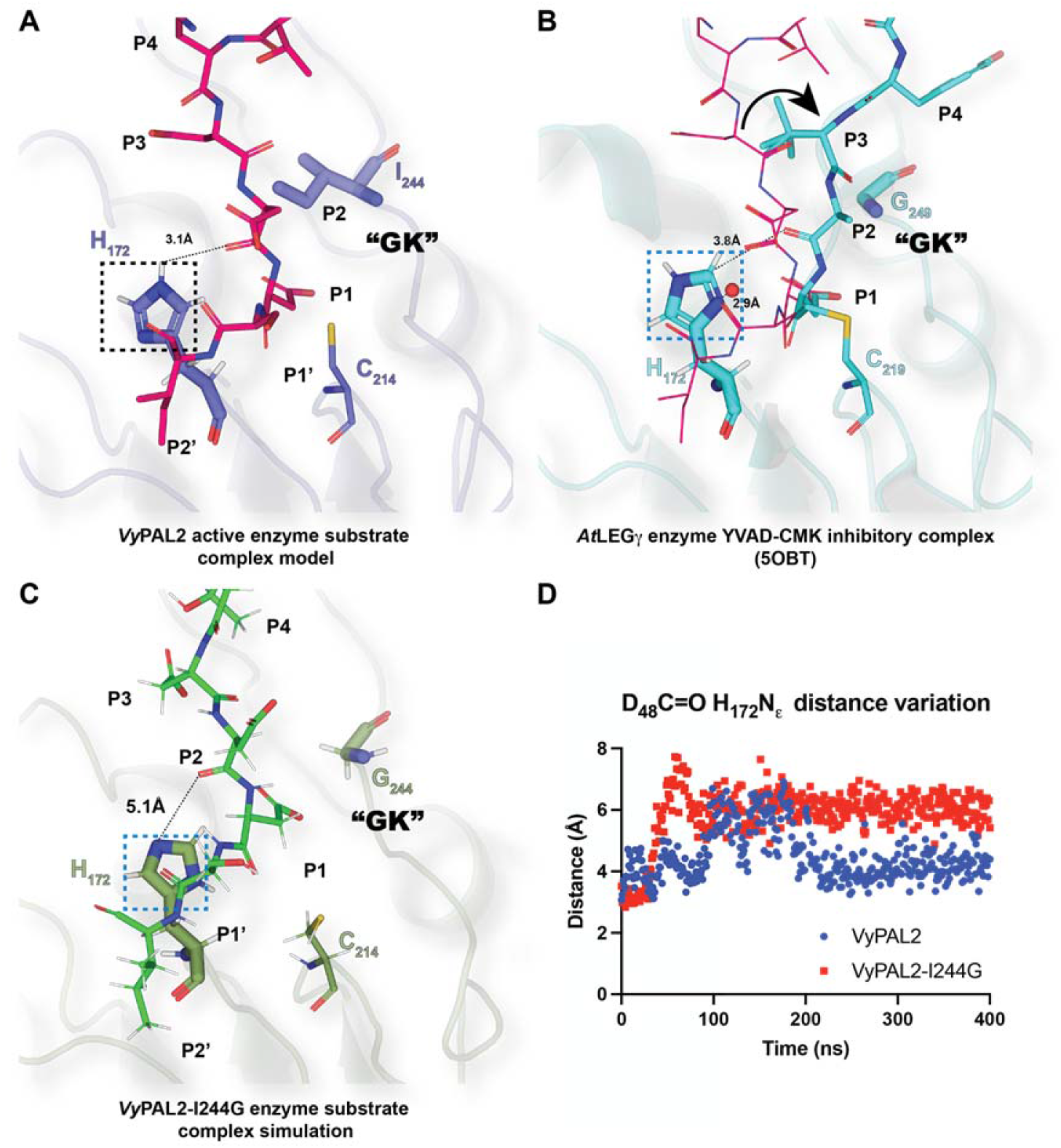
Role of the Gatekeeper position in substrate positioning. (A) Enzyme-substrate complex structure simulated based on the experimental VyPAL2-C214A structure (Supplemental Table 1, form I). Ala214 was replaced with a Cysteine. The enzyme core is shown as ribbons and colored in slate. The catalytic Cys214 and the His172 as well as the Ile244 gate-keeper residue are shown as sticks. The hydrogens of His172 are also shown. The P4-P2’ (NDDDSI) substrate is shown as sticks, colored in magenta and the 2Fo-Fc electronic density map contoured at 1σ and coloured in light blue. (B) Enzyme-inhibitor complex structure of AtLEGg (PDB code: 5OBT). The enzyme core is shown as ribbons and colored in cyan. The catalytic residues Cys219 and His172 as well as the Gly249 Gatekeeper residue are shown as sticks. The P4-P1 (YVAD-CMK) covalent inhibitor is shown as sticks, colored in cyan. The proposed nucleophilic water molecule shown as a red sphere is located at a distance of 2.9 Å from His172 in the axis of the thioester bond between the Cys219 and P1 Asp-CMK residue. For reference, the position of the VyPAL2-C214A substrate superimposed onto the AtLEGg bound structure is represented as lines and coloured in magenta. (C) Enzyme-substrate complex structure based on the VyPAL2-C214A-I244G MD simulation. The enzyme core is displayed as green ribbons. The catalytic Cys214 and His172 as well as the Gly244 gatekeeper residue are shown as sticks. The hydrogen atoms of His172 are also shown. The P4-P2’ (NDDDSI) substrate region is shown as sticks and coloured green. When the gatekeeper is a Glycine, the peptide is shifted and the distance between the carbonyl and the imidazole sidechain is 5.0 so that no hydrogen bond is formed. In this configuration, His172 is readily available to activate an incoming water molecule. (D) The distance between hydrogen atom of His172 Ne and the Asp48 carbonyl oxygen during the course the MD simulation is plotted.

The ligation reaction catalysed by VyPAL2 can be broken into two steps: firstly, a nucleophilic attack of the catalytic cysteine on the carbonyl of the P1 residue leading to acyl-enzyme formation breaking the peptide bond between the P1 and P1’ residues. This is followed in the second step by nucleophilic attack of the incoming peptide N-terminal amine on the acyl-enzyme and release from the catalytic cysteine. Here, the P1-P1’ scissile amide bond lies between the Asp49 and the Ser50 residues and the Asp49 carbonyl moiety is located at 2.9 Å from the Sγ atom of the catalytic Cys214 (Figure 3C). This distance is compatible with acyl-enzyme intermediate formation, confirming the pertinence of the present structure to represent an enzyme substrate complex.

### Detailed analysis of VyPAL2 substrate binding and specificity

The present structure is to our knowledge the first to disclose a complete enzyme-substrate complex for a peptide ligase allowing the complete definition of binding sites both at the prime and non-prime positions (Figure 4). It also allows the unambiguous assignment of residues belonging to the LADs: Here, LAD1 is formed by residues Trp243-Ile244-Thr245 and LAD2 comprises the Ala174-Pro175 dipeptide. LAD2 largely maps to the S2’ pocket albeit it does not form any direct interactions with the substrate P1’-P2’ residues. LAD1 impinges on the S2 and S3 pockets and Trp243 also contributes to shaping the S1 pocket (Figure 4).

Overall, the interface area between the substrate and the protein is 435 A^2^ and the total binding energy is -11.7 kcal/mol. As stated, P1 and P2’ residues constitute the main anchors to maintain the substrate with the largest buried surface areas upon complex formation of 112 Å^2^ and 128 Å^2^ in the S1 and S2’ pockets respectively (Figure 4). The S1 pocket is formed by negatively charged residues (at neutral pH) on one side of the pocket (Ser242, Asp263 and Glu212) and by positively charged residues on the other side (Arg69 and His70). This “polarity” favours tight binding of the P1 asparagine by providing hydrogen bonds to the oxygen and nitrogen atoms of its side chain. The carboxylate side chain of Asp49 closely overlaps with that of Asp of Ac-YVAD-CMK inhibitor in the bound AtLEGγ structure (PDB access code: 5OBT) and with the amide side chain of Asn from the AAN tripeptide in the bound HaAEP1 structure (PDB access code: 6AZT). At pH 4.5 of crystallization of the VyPAL2-C214A complex, Asp49 (P1 residue) is likely to be only partially negatively charged, while the sidechains of His70 and Arg69 are presumably positively charged. The negative charge carried by the carboxylate sidechain of Asp49 bound in the S1 pocket is neutralized by formation of two hydrogen bonds with the guanidinium group of Arg69 and with the imidazole group of His70 (Figure 4). Interestingly, space is left in the S1 pocket that can accommodate small adducts on the Asx sidechain, which agrees with the observation that N-hydroxy-Asn can also be used as a substrate by VyPAL2 (Xia et al., 2021).

On the primed side region of the substrate, in the S2’ pocket Tyr185 and the aliphatic chain of Lys177 establish van der Waals interactions with the P2’ residue. The floor of this pocket is formed by the backbone atoms of residues Ile178 and Gly179 at a distance of 3.6 Å from the Ile51 sidechain. The shape and chemical nature of this large pocket explains the selectivity for hydrophobic residues at P2’ observed in several PALs (Hemu et al., 2019; Nguyen et al., 2014). After P2’, the polypeptide chain of the substrate makes a sharp turn, such that no contact is made by P3’ residues and beyond (Figures 2 and 4).In contrast to P2’, the sidechain of Ser50 (P1’ residue) which has a smaller buried surface area of 72 Å^2^ (Figure 4), points away from the ligase molecular surface, which is consistent with the more relaxed specificity observed at P1’ position (Hemu et al., 2019; Nguyen et al., 2014). the P1’ residue appears to play an indirect role for binding: while a Pro residue would induce a rigid kink in the peptide structure and is therefore disfavoured, the presence of a Gly residue at P1’ is likely to endow the peptide with the flexibility required for a deep anchoring of the neighbouring hydrophobic P2’ residue (Figure 4). Thus, a variety of residues such as His, Lys, Asn, Gln, Arg or Ser can be accommodated at P1’ with good cyclization activity (Hemu et al., 2019). Conversely, residues such as Ile, Val, Tyr or Trp might introduce non-favourable hydrophobic interactions with mostly the polar surface of the VyPAL2 ligase at this location.

On the non-prime side, The P2 Asp48 and P3 Asp47 residues have respective buried surface areas of 83 Å^2^ and 72 Å^2^ upon complex formation. The sidechain of Asp48 (P2) also points away from the ligase binding groove suggesting that many residues can be accommodated at this position (Figure 4), while the carboxylate group of Asp47 (P3) makes a salt-bridge with the guanidinium group of Arg69 (Figure 4). Asp48 at the P2 position has its backbone carbonyl involved in a hydrogen bond with the imidazole group of the catalytic His172. It also interacts with the P1’ residue of the substrate via a hydrogen bond between the respective side chains of Asp48 and Ser50.

In summary, the pattern of recognition revealed by the present complex suggests only two main anchoring residues at P1 and P2’ with restricted specificity, while P1’ and P3 are amenable to more sequence variations, and no sequence restrictions for P2 and residues beyond P3’. It largely explains the efficiency with which Asn ligase such as butelase 1 or VyPAL2 can accommodate a large variety of bulky protein substrates as ligands and can even perform protein-protein ligation or protein cyclization.

### The “Gatekeeper” residue induces a shift in the substrate position at the non-prime site

It was previously observed that mutation of a Cys residue (Cys247) near the active site of OaAEP1b (PDB access code: 5H0I), to larger amino acids (Thr, Met, Val, Leu, Ile), reduced ligation catalytic efficiency, while mutations to smaller residues such as Ala resulted in over a hundred-fold improved ligation efficiency. Moreover, mutation of this “Gatekeeper” residue into Gly results in an increased amount of hydrolysis product, suggesting that residues at this site play a major role in modulating enzyme function(Yang et al., 2017). Interestingly, MD simulation suggested a shift in the position of the N-terminal (non-prime) portion of the substrate due to steric hindrance introduced by a gatekeeper residue bulging at the surface of the S2 substrate-binding pocket (Hemu et al., 2019). Here, the experimental structure of the ligase-peptide complex confirms and extends this hypothesis: while there is excellent substrate overlap at the P1 position between VyPAL2-C214A form I (this work) and AtLEGγ covalently bound to Ac-YVAD-cmk (PDB access code: 5OBT), a repositioning of residues P2-P3 of the substrate occurs in the presence of a protruding sidechain at the Gatekeeper position (Figure 5; Supplemental Figure 6): In the presence of Ile244 as gatekeeper, P2-P3 residues are pushed away by about 4 Å from the ligase active site surface compared to the position adopted when the gatekeeper is a Gly, allowing the peptide substrate to come closer to the enzyme surface. As a result, a hydrogen bond forms between the backbone carbonyl oxygen of the substrate P2 residue (Asp48) and the imidazole ring of catalytic His172 (Figure 5A). This hydrogen bond is possible in the orientation of the histidine imidazole ring adopted in the experimental structure with the Nε atom oriented towards the substrate. Conversely, reorientation of the histidine sidechain towards the catalytic cysteine, as seen in AEP structures, enables the activation of an attacking water molecule (Figure 5B; Supplemental Figure 6). The relevance of these observations for ligation activity was then tested using MD simulations.

**Figure 6.**
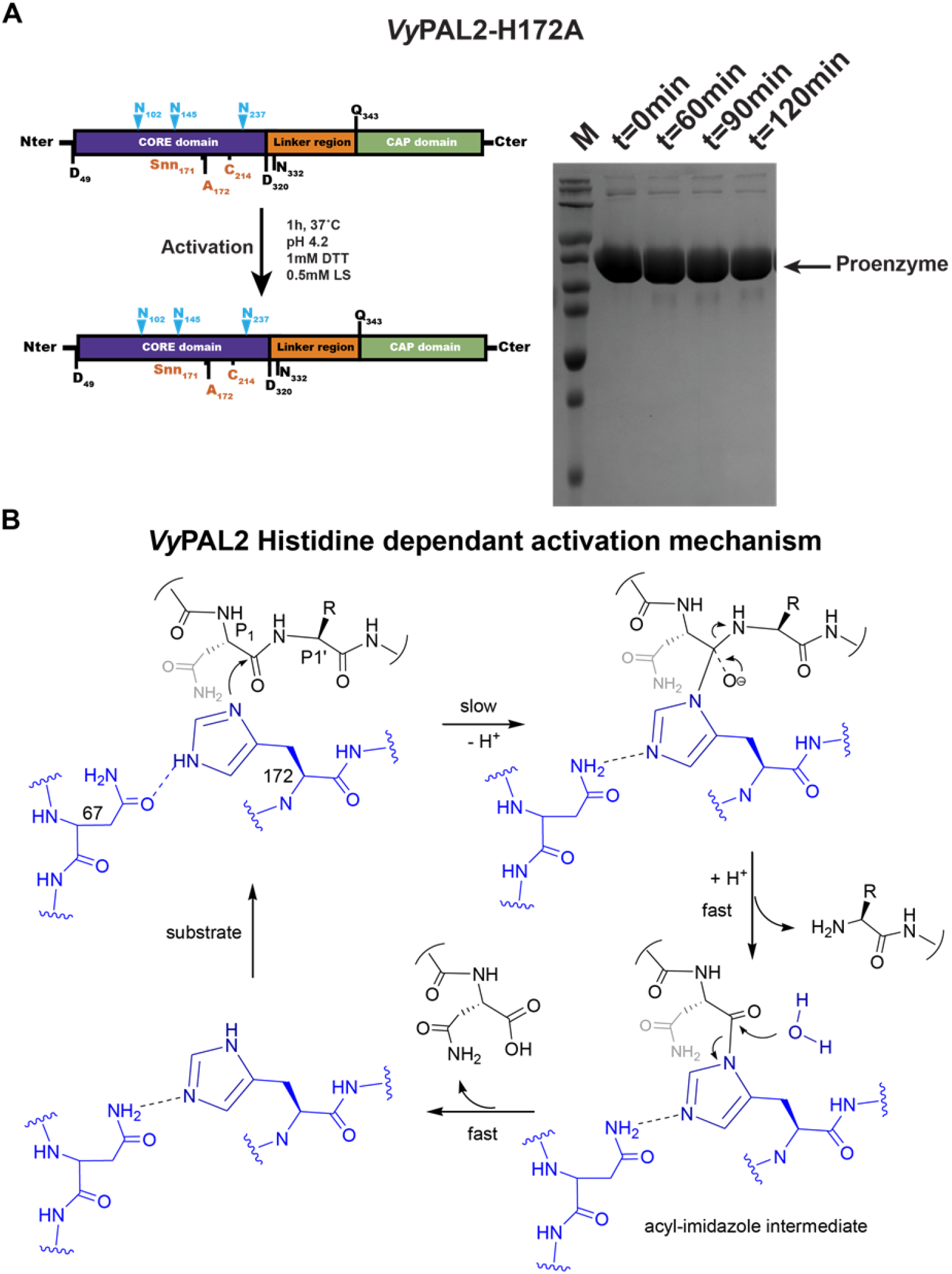
His172 is the catalytic residue necessary for proenzyme VyPAL2 activation. (A) Schematic sequence representations of VyPAL2-H172A. The products from low pH processing of the proenzyme were analysed using SDS-PAGE (right panel) with Coomassie blue staining. Incubation for up to 2 hours of VyPAL2-H172A did not lead to any processed enzyme. (B) Proposed mechanism of processing of VyPAL2 that is Cys214 independent and where the His172 is the main catalytic residue. This mechanism involves the formation of an acyl-imidazole intermediate that is highly unstable and rapidly resolved by the attack of a nucleophilic water molecule.

We submitted two enzyme-substrate complexes to Molecular Dynamics simulations. The enzyme substrate complexes were formed using residues 53-328 of the VyPAL2-C214A structure as the enzyme and residues 43-51 from the neighbouring asymmetric molecule as substrate. A second model, where the residue Ile244 was computationally replaced by a Gly244, was similarly produced. Both enzyme-substrate complexes were simulated for 10 ns. In the MD simulation the most noticeable consequence of the I224G mutation is a flip of the catalytic His172 side chain imidazole ring that occurs rapidly (after 1 ns of simulation) (Figure 5C). This movement leads to the catalytic histidine side chain to adopt the conformation observed in AEP structures for the rest of the simulation (Figure 5D). As a consequence of this flip, the hydrogen bond between the carbonyl oxygen of the substrate P2 residue (Asp48) and the imidazole ring of catalytic His172 is lost. The distance between the two moieties rapidly increases above 5 Å and is stabilized at this level in the case of the VyPAL2-I244G mutant (Figure 5D). This is in contrast with the MD simulation of the Ile244 structure, which shows very little change for the entirety of the simulation. Both the histidine sidechain conformation and Asp48 carbonyl distance to the Nε, are stable, allowing for stable hydrogen bond formation. In summary, the presence of a Gly residue at the gatekeeper position is conducive to a positioning of the catalytic His such that it can easily activate an incoming water molecule for hydrolysis, while a bulkier residue such as Ile leads to a hydrogen bond formation of the same histidine with the substrate that is detrimental to hydrolysis and favours ligation.

### VyPAL2 can be auto-activated solely by its catalytic histidine

PALs and AEPs are naturally expressed as proenzymes that require activation to expose their active site to protein and peptide substrates. The activation steps consist of sequential hydrolysis events targeting an N-terminal pro-peptide of ∼50 residues and the C-terminal cap domain (∼150 residues). Auto-activation of AEPs and PALs represents a proteolysis reaction regardless of the main activity (protease or ligase) of the mature enzyme. It is commonly thought that the auto-activation mechanism involves the nucleophilic attack of the catalytic cysteine onto the peptide carbonyl at Asn and/or Asp residues in the N-terminal and the linker regions of the pro-enzyme. In this light, the ability of VyPAL2-C214A to efficiently auto-activate at acidic pH was rather surprising as it lacks the catalytic cysteine to perform the nucleophilic attack. To further explore the auto-activation mechanism, we generated a single mutant of VyPAL2 having its catalytic Histidine mutated to Alanine. The VyPAL2-H172A mutant was unable to auto-activate (Figure 6A) as no lower apparent molecular weight band was observed and no reduction in the pro-enzyme occurred during the assay. This result could suggest an alternative route for VyPAL2 auto-activation in the absence of a catalytic Cys214. This route would rely solely on His172 and would not be linked to the mature protein activity be it ligation of hydrolysis. Indeed, we did not observe any significant activity for the processed VyPAL2-C214A mutant in a hydrolysis or ligation assays at either neutral or acidic pH.

Upon acidification, numerous interactions between charged residues at the interface between the core and cap domains are broken, leading to an intermediate structure having two domains connected by the linker region (Movie S1). This is followed by proteolysis at several Asx residues, easily accessible in the linker domain. Here, in the case of VyPAL2, three major activation sites were found in the N-terminal propeptide and one in the linker region (Figures 1 and 2). Our results show that partial enzyme activation can occur without formation of an S-acyl-enzyme intermediate: the VyPAL2-C214A mutant can rapidly cleave the cap domain while complete processing including proteolysis of the N-terminal pro-peptide only happens after weeks of incubation (Figure 1 and form II VyPAL2-C214A crystals). A similar observation was previously done for a legumain having its catalytic Cys mutated to Ala, which was also able to undergo activation (Zhao et al., 2014). Intriguingly, although the VyPAL2-C214A mutant can very efficiently perform auto-proteolysis, it does not show any hydrolytic activity at acidic pH using a standard peptide (GN14: GISTKSIPPISYRNSL (Hemu et al., 2019)) as substrate, an observation which also holds true for VyPAL2. In contrast, the presence of the catalytic His172 is absolutely required for auto-activation, since VyPAL2-H172A cannot be activated.

## DISCUSSION

Proteases and ligases share a high amino-acid sequence and structural identity. PALs could have divergently evolved from AEPs to catalyse transpeptidation or ligation reactions required to produce cyclic peptides in plants as a mechanism for defence against pathogens. At the molecular level, the nature of this evolution was recently clarified with the characterisation of the Gatekeeper residue and Ligase Activity Determinants (LADs) concepts. This notion proposes that only a few specific positions in the substrate binding site are sufficient to convert an AEP into a PAL. In this work we have uncovered a possible molecular mechanism accounting for the primary role of the Gatekeeper residue in determining the direction of the enzyme activity.

Ligation or hydrolysis reactions appear to proceed through two shared initial steps: (i) substrate binding and nucleophilic attack by the Cys sulfhydryl moiety on the P1 carbonyl (Figure 3C) leads to the formation of an acyl-enzyme intermediate and cleavage of the P1-P1’ peptide bond (ii) nucleophilic attack onto this acyl-enzyme intermediate breaks the transient thioester bond formed between the catalytic cysteine and the P1 Asx residue. The main difference between ligation and hydrolysis derives from the selection of the incoming nucleophilic group to resolve the acyl-enzyme intermediate: either a water molecule for proteolysis or an amine from an incoming peptide for ligation. This proposed overall scheme concurs with the elegant D_2_O exchange measurements that demonstrated the absence of an isotopic shift in a kalata B1 cyclotide peptide produced by peptide cyclization reaction, hence ruling out the role of a water molecule in the ligation reaction carried by OaAEP1b (Harris et al., 2015) and favouring the scheme where direct aminolysis and transpeptidation is performed by an incoming amine (Hemu et al., 2019).

PALs and AEPs share almost identical structures and subtle differences at their LAD1 and LAD2 regions play a key role in determining the direction of the reaction (Hemu et al., 2019). The LAD2 region, located at the primed side of the peptide binding site was shown to alter the enzyme preference: non-hydrophobic residues at LAD2 favour the presence of water molecules while bulky residues tend to exclude an incoming peptide. For instance, the Y168A mutation targeting LAD2 was sufficient to endow the VyPAL3 and VcAEP proteases with significant peptide cyclase activity18. Conversely, a small exposed hydrophobic residue like Alanine at this position will favour ligation as its hydrophobicity would decrease the local affinity for water molecules from the solvent, while its small size can easily accommodate an incoming peptide (Supplemental Figure 7). Here, as seen in the form I complex, VyPAL2 possesses Ala174-Pro175-Gly176 at the LAD2 region which both maintain the local secondary structure of the protein (a β-turn) and have the required size and hydrophobicity to bind Ile51 as P2’ of an incoming peptide.

Another key determinant of activity lies in the Gatekeeper residue centred at LAD1 on the non-prime side of the binding site. In the VyPAL2-C214A form I complex, the Gatekeeper (Ile244) and the catalytic His172 are located at opposite sides of the peptide binding groove. The aliphatic sidechain of Ile244 lies at van der Waals contact distance of 3.3 Å from the α-carbon residue P2 (Asp48) of the peptide ligand (Figure 3). A comparison with enzymes having a Gly residue as Gatekeeper such as AtLEGγ covalently bound to Ac-YVAD-cmk, reveals that the corresponding P2-P3 residues are displaced by about 4 Å from the ligase active site (Figure 5; Supplemental Figure 6). As a result, the displacement induced by an aliphatic sidechain as Gatekeeper, allows the formation of a hydrogen bond between the carbonyl oxygen of the substrate P2 residue (Asp48) and the imidazole ring of catalytic His172 (Figure 5). In contrast, when the Gatekeeper is a glycine, the corresponding distance is too far to establish a hydrogen bond (Figures 5B to 5D). Accordingly, this hydrogen bond is not observed in other active form protease structures. Thus, the present orientation of the imidazole ring of His172, constrained by this polar interaction with the peptide, seems incompatible with a role in activating an incoming water molecule for nucleophilic attack and appears to be a unique feature of a ligase (Movie S2). Conversely, in AEPs, the catalytic histidine can act as a base to assist in the activation of the water molecule that is positioned above Gly173 in VyPAL2, (corresponding to Gly178 in the case of AtLEGγ for the nucleophilic attack of the thioester bond. We propose that this subtle change in the peptide substrate conformation induced by the sidechain of the Gatekeeper explains why PALs favour a nucleophilic attack by the amine group of an incoming peptide, and why at a more acidic pH, the ligation reaction becomes less favoured due to protonation of the incoming amine group.

Our results have shown that the auto-activation can be performed without the presence of the catalytic Cys, while the catalytic His is necessary (Figures 2A and 6A), suggesting that a different molecular mechanism is at play between auto-activation and peptide hydrolysis/ligation. Based on our results, we proposed a mechanism for the auto-activation of VyPAL2-C214A. At the activation pH, His172 and Asn67 likely act as catalytic dyad. The imidazole nitrogen of His172 acts as nucleophile to attack and cleave the P1-P1’ scissile bond, forming an acyl-imidazole intermediate. As protons are continuously taken up and given back to surrounding solvent water molecules near the catalytic site, once formed, the acyl-imidazole intermediate could be quickly hydrolysed (Figure 6B). Indeed, at the acidic pH of 4-5, the imidazole of catalytic histidine residue can act as both an acid and a base. Therefore, it is conceivable that one of the nitrogens on the imidazole of His172 in VyPAL2-C214A mutant can serve as a nucleophile to attack the carbonyl carbon of the scissile peptide bond, forming an acyl-imidazole intermediate which is very unstable and undergoes rapid hydrolysis. The amide hydrogens of Gly173 and Ala214 form hydrogen bonds with the carbonyl oxygen of the scissile peptide bond. The same hydrogen bonds can still form with the oxyanion for its stabilization, which is crucial for the formation of the acyl-enzyme intermediate because the first step of this mechanism has the highest activation energy and hence constitutes the rate-determining step.

## CONCLUSION

In summary, we found that the N-terminus of VyPAL2-C214A containing the Asp49-Ser50-Ile51 sequence constitutes a P1P1’P2’ substrate proteolytic motif allowing us to perform a detailed structural analysis of the enzyme active site pockets that are used for both protease and ligase activities. In this respect, peptide asparaginyl ligases appear as opportunistic catalysts that have evolved from asparaginyl endoproteases: by recycling essentially the same binding site with the introduction of only a few key residues near the conserved S1 pocket. A slight distortion induced in the substrate peptide conformation appears sufficient to endow the same catalytic site with a novel activity. This dual usage of the specificity pockets is also reflected in the way we can interpret the present form I crystal structure: the complex can be seen either as a pro-enzyme-substrate complex or alternatively as a ligase-product complex. In the former, the proenzyme is caught in the act of cleaving in trans the N-terminal tail of a companion molecule for maturation, whilst in the latter, the ligase has just performed a protein-peptide ligation reaction using Asp as P1 residue.

## METHODS

### Protein expression and auto-activation

The expression and purification of VyPAL2 and the different mutants were carried out as previously described (Hemu et al., 2019). Briefly after mutagenesis, the genes coding for VyPAL and its mutants were expressed in Sf9 insect cells using the Bac-to-Bac protocol (Invitrogen). Protein purification was performed in three steps with IMAC affinity purification followed by ion-exchange and size-exclusion chromatography (SEC, with 1x PBS, pH 7.4, 1 mM DTT). The protein was then concentrated and stored at 4 °C.

Following gel filtration, proenzymes were concentrated to 2 mg/ml. The optimal activation time was estimated following a time-course analysis. Zymogens were activated at 37 °C in 50 mM citrate, 100 mM NaCl in a 1:1 volume ratio (v/v), with addition of 0.5 mM N-Laurocryosine, 1 mM DTT, final pH 4.4. The samples were checked using SDS-PAGE to determine the optimal activation time. After scaled-up proenzyme activation, the activation mixture was subjected to an SEC purification with buffer containing 20 mM MES, pH 6.5, 0.1 M NaCl, 1 mM DTT. Fractions containing the active form protein were pooled and concentrated for further use.

### Crystallization, Data Collection, and Structure Determination

The active enzymes were concentrated to 4∼9 mg/ml and screened against JCSG-plus HT-96 (Molecular Dimensions) screening kit. Crystals were obtained for the active form of VyPAL2 (6.7 mg/ml): 0.1 M sodium acetate, pH 4, 0.2 M lithium sulfate, 30% PEG 8000; VyPAL2-I244V (10.4 mg/ml): 0.1 m Bis-Tris pH=7.5, 25% (w/v) PEG 3350; VyPAL2-C214A (Form I, 4.93 mg/ml): 0.2 M Lithium sulfate, 0.1 M sodium acetate, 30% (w/v) PEG 8000 pH=4.6 and for VyPAL2-C214A (Form II, 9.14 mg/ml): 0.2 M potassium formate, 20% (w/v) PEG 3350. Crystals suitable for X-ray data collection were frozen in liquid nitrogen with addition of 20% (w/v) glycerol and shipped to synchrotrons for data collection. Data were integrated using XDS (Kabsch, 2010) and the structures were solved using 6IDV as a model in Phaser (CCP4) (Winn et al., 2011). Manual corrections of the structures were performed in the Coot program for molecular graphics (Emsley et al., 2010) and refined using Buster TNT (GlobalPhasing Ltd) (Smart et al., 2012). Processing and refinement statistics are presented in Supplemental Table 1. The structures were deposited in the Protein Data Bank.

### Kinetic studies

Substrate peptides were synthesized by Bio Basic and consist of an N-terminal substrate PIE(EDANS)YNAL and C-terminal substrate GIK(DABSYL)SIP. Ligation assays were performed in 100 µL reaction mixture containing 60 µL of MES buffer (20 mM MES, pH 6.5, 0.1 M NaCl, 1 mM DTT), 20 µL of serial dilutions (with the MES buffer) of substrates containing both N- and C-terminal substrates with ratio (EDANS: DABSYL) of 1: 3. The final concentrations of EDANS substrate were 50 µM, 25 µM, 12.5 µM, 6.25 µM, 3.125 µM, 1.5625 µM, and 0.78125 µM. The enzyme was injected right before measurement by the injector of Tecan 10M Spark microplate reader; 20 µL of the enzyme at 100 nM was added to give a final concentration of 20 nM. The reactions were run at 37 °C for 2 min and measured at 5-second intervals to calculate the initial rates during the linear portion of the progress curve. All samples were added to Thermo Fisher Scientific-Nunclon 96 Flat Black plate, and fluorescence was measured using Tecan 10M Spark microplate reader with the excitation wavelength at 336 nm and emission wavelength at 490 nm.

### Molecular Dynamics simulation

To obtain the equilibrated position of the substrate, the enzyme-substrate complexes were subjected to Molecular Dynamics simulations on NAMD 2.12.3 (Phillips et al., 2016). The complexes are formed by using the 43-51 residues from the neighbouring as the substrate and the residues 52-326 as the core domain. The complexes were simulated in a water box where the minimal distance between the solute and the box boundary was 15 Å along all three axes. The charges of the solvated system were neutralized with counter-ions, and the ionic strength of the solvent was set to 150 mM NaCl using VMD.4 (Humphrey et al., 1996). The fully solvated system was subjected to conjugate gradient minimization for 10,000 steps, subsequently heated to 310 K in steps of 5 ps. The system was simulated for 20 ns with the Ca atoms of Asp49 of the peptide substrate fixed in space and the backbone atoms of the ligase constrained by a harmonic potential of the form U(x)=k(x-xref)2, where k is 1 kcal mol-1 Å-2 and xref is the initial atom coordinates. Additionally, the distance between the Sγ atom of Cys214 and the main-chain C atom of Asp49 are kept at 2.94 Å under a harmonic potential. Such constraints allow the side chains of enzymes and the rest of the peptide substrate to move freely. All simulations were performed under the NPT ensemble assuming the CHARMM36 force field for the protein5 and assuming the TIP3P model for water molecules (Best et al., 2012). The MD simulations were performed on ASPIRE-1 of the National Supercomputing Centre, Singapore (https://www.nscc.sg).

## Supplemental Data

**Supplemental Table 1**. Data collection and refinement statistics.

**Supplemental Table 2** R.M.S.D. of *Vy*PAL2 with other plant AEPs (PDBeFold).

**Supplemental Figure 1**. SEC profiles of VyPAL2 wt and I244V.

**Supplemental Figure 2**. Electrostatic charges at the molecular interface of the mature *Vy*PAL2 active domain and cap region.

**Supplemental Figure 3**. Activation kinetics comparison between VyPAL2 and VyPAL2-C214A.

**Supplemental Figure 4**. Comparison of the active structures of VyPAL2 and its mutants.

**Supplemental Figure 5**. Position of proenzyme cap domain Gln343 and VyPAL2-C214 active form Asp49.

**Supplemental Figure 6**. Effect of LAD1 “Gate keeper” residue on the molecular surface of *Vy*PAL2.

**Supplemental Figure 7**. Effect of the presence of large bulky residue a the LAD2 region.

**Supplemental Movie 1**. Activation of VyPAL2.

**Supplemental Movie 2**. Structure comparison between VyPAL2-C214A Form I and AtLEGγ bound to Ac-YVAD-cmk.

## ACKNOWLEDGMENTS

This research was supported by Academic Research Grant Tier 3 (MOE2016-T3-1-003) from Singapore Ministry of Education (MOE) to the JL, JPT and CFL laboratories. We thank for their expert assistance scientists and beamline staff at (i) Spring-8 Synchrotron where VyPAL2 active enzyme structure was collected, (ii) Swiss Light Source (SLS) where VyPAL2-C214A form II was collected, and (iii) MX2 (Australian Synchrotron) where VyPAL2-C214A form I and VyPAL2-I244V datasets were collected. We thank the National Supercomputing Centre, Singapore (https://www.nscc.sg) for providing ASPIRE-1 for the MD simulations.

